# Enzymatic cleaving of entangled DNA rings drives scale-dependent rheological trajectories

**DOI:** 10.1101/2023.12.05.570275

**Authors:** Philip Neill, Natalie Crist, Ryan McGorty, Rae Robertson-Anderson

## Abstract

DNA, which naturally occurs in linear, ring, and supercoiled topologies, frequently undergoes enzyme-driven topological conversion and fragmentation in vivo, enabling it to perform a variety of functions within the cell. In vitro, highly concentrated DNA polymers form entanglements that yield viscoelastic properties dependent on the topologies and lengths of the DNA. Enzyme-driven alterations of DNA size and shape therefore offer a means of designing active materials with programmable viscoelastic properties. Here, we incorporate multi-site restriction endonucleases into dense DNA solutions to linearize and fragment circular DNA molecules. We pair optical tweezers microrheology with differential dynamic microscopy and single-molecule tracking to measure the linear and nonlinear viscoelastic response and transport properties of entangled DNA solutions over a wide range of spatiotemporal scales throughout the course of enzymatic digestion. We show that, at short timescales, relative to the relaxation timescales of the polymers, digestion of these ‘topologically-active’ fluids initially causes an increase in elasticity and relaxation times followed by a gradual decrease. Conversely, for long timescales, linear viscoelastic moduli exhibit signatures of increasing elasticity, and DNA diffusion, likewise, becomes increasingly slowed —in direct opposition to the short-time behavior. We argue that this scale-dependent rheology arises from the population of small DNA fragments, which increases as digestion proceeds, driving self-association of larger fragments via depletion interactions, giving rise to slow relaxation modes of clusters of entangled chains, interspersed among shorter unentangled fragments. While these slow modes dominate at long times, they are frozen out in the short-time limit, which instead probes the faster relaxation modes of the unentangled population.

## Introduction

The cellular environment exists in a state far from equilibrium – a necessity to perform complex cellular functions ^1^. Densely crowded biopolymers such as proteins and DNA are constantly manipulated by molecular motors and enzymes which drive changes to the architecture and transport of the biopolymers, which in turn alter the viscoelastic properties of the cellular environment ^2–6^.

Much work over the last two decades has been devoted to how the action of molecular motors (e.g., myosin, kinesin) on cytoskeleton filaments (e.g., actin, microtubules) drive mechanical changes to in vitro cytoskeletal networks, as well as the cellular environment ^7–11^. In these systems, the motors consume energy to actively pull on the filaments to drive mechanical changes.

However, only recently has the action of restriction enzymes, that alter the topology and/or length of biopolymers such as DNA, been appreciated as a potential driver of changes to viscoelastic properties ^12–14^. Yet, this process is ubiquitous in diverse processes including nuclear division, DNA repair, transcription and signaling ^1,3,15,16^. For example, DNA naturally occurs in supercoiled circular, relaxed circular (ring), and linear topologies with lengths that scale many decades. A host of restriction endonucleases naturally exist to cleave DNA strands at specific sites, via basepair recognition, to convert supercoiled or ring DNA to linear form, with a single cut, and subsequently cleave the linear strand into smaller fragments if multiple recognition sites exist along the chain ^15^.

Recent work demonstrated that topology-altering enzymatic digestion of DNA can push concentrated DNA solutions out of equilibrium to elicit time-varying microrheological properties of the solutions en route from their initial, undigested steady-state to their fully-digested steady-state ^13^. In these studies, the viscosity of ‘topologicallyactive’ solutions of 5.9 kilobasepair (kbp) DNA was shown to steadily increase under the action of a single-site restriction enzyme that cleaves each circular DNA molecule into a linear strand of the same length. The degree to which the viscosity increased during linearization depended on the enzyme:DNA stoichiometry as well as the concentration of the DNA. This effect was shown to arise from the changing degree of polymer overlap as rings are converted to linear chains. Namely, the radius of gyration of a circular or ring polymer is smaller than that of its linear counterpart, following the relation *R*_*G,L*_ ≃ 1.58*R*_*G,R*_ ^17^; and because the polymer overlap concentration 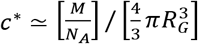 scales as 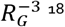 ^18^, as circular DNA is converted to linear topology, the degree of polymer overlap *c/c*^*^ increases despite the fact that *c* does not change.

This same study showed that fragmentation of entangled linear DNA caused the viscosity to steadily decrease, as expected as the degree of polymer overlap also decreases with decreasing polymer length *L* ^13^. Specifically, given the molecular weight of a polymer scales with its length *M*∼*L* and for an ideal random coil *R*_*G*_∼*L*^1*/*2^ it follows that *c*^*^∼*L*^−1*/*2^ such that *c/c*^*^∼*L*^1*/*2 19^. The dependence of overlap and entanglements on length can also be seen in the reptation model relation between polymer length *L* and the polymer length between entanglements *l*_*e*_∼*c*^−1.31 18^. Because *l*_*e*_ is set by the concentration, the shorter the chains are at a given concentration the fewer entanglements per chain (*L/l*_*e*_), which dictate the viscoelastic response for entangled polymers.

Following on this work, bulk rheology studies of topologically-active 11-kbp ring DNA undergoing digestion by a 13-site cutter showed an unexpected sharp transition from a low-elasticity state to a high-elasticity state, marked by a ∼50-fold increase in the elastic modulus *G*′, upon complete digestion ^14^. Not only is the rise in elasticity at odds with the decreasing viscosity measured for fragmenting linear chains described above, but the transition between the states was also not steady or continuous, as was reported for microrheology measurements ^13^. This same study also showed that the same enzymatic activity elicited very different rheological trajectories for composites of DNA and dextran with varying volume fractions of DNA. Namely, while all systems displayed the initial abrupt jump from low-elasticity to high-elasticity, this transition occurred much earlier in the digestion process for DNA-dextran composites, as compared to 100% DNA, and was followed by a subsequent drop back to a low-elasticity state. These results were shown to arise from cooperative percolation of clusters of slow (long, entangled, linear) DNA, facilitated via crowding by small dextran polymers, and that dictate the rheological response ^14,20–22^.

The studies described above highlight the different responses that topologically-active DNA systems can exhibit at different spatiotemporal scales, and the important role that microscale heterogeneities, crowding, and depletion interactions may play. Moreover, the effect of end closure of ring polymers on their ability to form entanglements and elicit elastic-like response, a topic of intense interest in the polymer community ^20,23–29^, also contributes nontrivially to the rheological trajectory of these topologically active systems. In particular, linearization of rings is expected to increase the elastic contribution to the stress response; however, threading of rings by linear chains in transient ring-linear blends may exhibit even stronger constraints and slowed relaxation than transient 100% linear solutions ^30–37^.

While the microscale and macroscale responses of topologically-active fluids are clearly different, the elusive mesoscale rheology that connects these two limits remains an important unknown. How the time-varying topologies of the molecules effect their response to nonlinear straining, at rates much faster than the intrinsic relaxation rates of the system, is also an essential unanswered question. Finally, how the diffusion of the molecules undergoing digestion varies to give rise to the non-equilibrium rheological response is critical to mapping the dynamics across spatiotemporal scales.

Here, we design topologically-active solutions of circular PYES2 DNA of length *L* = 5.9 kilobasepairs (kbp) at a concentration well above the overlap concentration of the initially circular DNA, 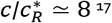 ^17^. We note that when the solution is transiently fully linearized (but not yet fragmented), at the beginning of the digestion, the degree of overlap is ∼4x higher 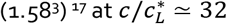. We incorporate a multi-site restriction endonuclease HaeIII, that cleaves pYES2 at 17 different recognition sites, creating 17 blunt-ended linear strands ranging in length from 19 bp to 1692 bp (Figs 1A, S1). To address the issues raised above, we measure the time-resolved rheological properties of the DNA solutions across three decades of spatiotemporal scales from *t* ≃ 17 ms to *t* ≃ 6.3 s and from *x* ≃ 20 nm to *x* ≃30 μm. We achieve this breadth by combining both linear and nonlinear optical tweezers microheology measurements (OTM) (Fig 1B-F) with differential dynamic microscopy (DDM) analysis of DNA fluctuations (Fig 1G-J).

**Figure 1:**
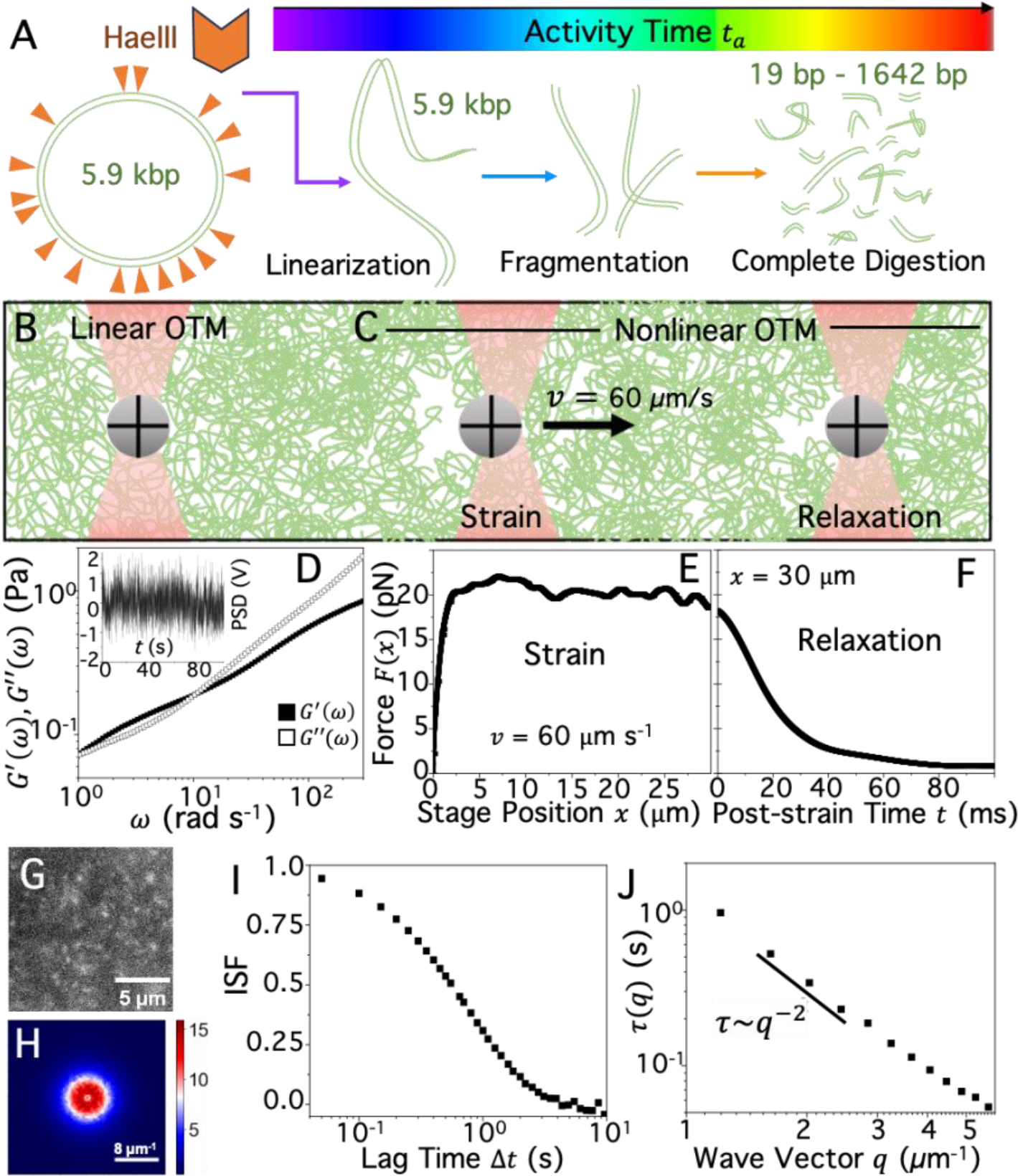
Optical tweezers microrheology and differential dynamic microscopy (DDM) couple time-dependent rheology with transport properties of topologically-active DNA across spatiotemporal scales. **(A)** Digestion of 5.9 kbp plasmid DNA (pYES2) by restriction enzyme HaeIII first linearizes the initially circular DNA molecules then successively cleaves the full-length linear chain into shorter fragments over activity time *t*_*a*_, resulting in 17 linear fragments of lengths *l*_*f*_ = 19 bp to 1642 bp at the end of digestion. **(B**,**C)** Cartoon of optical tweezers microrheology (OTM) probing both the linear (B) and nonlinear (C) force response of concentrated DNA solutions as they undergo HaeIII-driven topological digestion. (B) A 4.5 μm microsphere probe (grey) embedded in a DNA solution (green) and trapped by an infrared laser (red). The thermal fluctuations of the trapped probe from the surrounding DNA solution are measured by recording the corresponding deflections of the trapping laser via a position sensing detector (shown in inset in (D)). **(C)** The same microsphere is dragged through the DNA solution at a constant speed of *v* = 60 μm/s (left) through a distance *s* = 30 μm, using a nanopositioning stage to move the sample relative to the trap. After this displacement, the stage motion is halted, and relaxation of the force on the probe from the surrounding DNA relaxing strain-induced stress is measured (right). (**D**) Thermal fluctuations measured at 20 kHz over the course of 100 s (shown in inset) are used in linear OTM to compute the frequency-dependent linear viscoelastic moduli, *G* ^′^(*ω*) (filled squares) and *G* ^′′^(*ω*) (open squares), using the generalized Stokes-Einstein relation (GSER), as described in Methods. **(E**,**F)** In nonlinear OTM, the force exerted on the probe by the DNA is measured during (E) and following (F) the 30 μm displacement to determine the strain response and relaxation dynamics. **(G)** To measure DNA diffusion using DDM, a small fraction of DNA is fluorescent-labeled and time-series of the diffusing molecules are recorded. **(H)** DDM analysis results in an image structure function 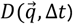 as a function of lag time Δ*t* and wave vector *q* in Fourier space. Shown is 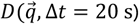. The radially symmetric image structure function is azimuthally averaged to arrive at a one-dimensional function *D*(*q*, Δ*t*) from which we extract the intermediate scattering function (ISF). **(I)** Sample ISF as a function of lag time Δ*t* for *q* = 2 μ**m**^−1^, which is fit to an exponential to determine the (**J**) density fluctuation decay time τ which exhibits power-law scaling τ(*q*)∼*q*^−2^, indicative of diffusive dynamics.

We perform measurements for two different HaeIII:DNA stoichiometries of 0.3 U/μg and 0.5 U/μg, chosen based on previous results, to be high enough to achieve complete digestion over the course of <4 hours but low enough such that the system can be considered in quasi-steady state during each individual measurement. Specifically, the longest measurement timescale we probe (<10 s) is several orders of magnitude shorter than the time to completely digest the DNA, *t*_*d*_ ≈6000 *s*, and time for a single cut *t*_*d*,1_ ≈*t*_*d*_*/*17 ≃ 350 *s*. We also note that there is no noticeable change in the topology of the molecules in the solution over intervals shorter than ∼5 mins ( 300 s), consistent with *t*_*d*,1_. This separation of timescales, between measurement times *t* and enzymatic activity time *t*_*a*_ = 10 – 240 min, allows us to treat the solutions during each measurement as in quasi-steady state, such that we can evaluate steadystate quantities such as viscoelastic moduli, *G*′(*ω*) and *G*′′(*ω*), and diffusion coefficients *D*. Throughout the paper, we will refer to the time associated with digestion as activity time *t*_*a*_, the variation in measured quantities as a function of *t*_*a*_ as trajectories, and the timescales probed in each measurement as the measurement timescale or time *t*.

Our results uncover stark differences between the trajectories of rheological metrics that probe short versus long measurement timescales. Namely, the response of topologically-active solutions to fast strain rates (short times) appear to first increase in elastic-like contributions follows by a large drop before reaching steady-state force and viscosity values that are substantially lower than the solution before digestion. In direct opposition, the slow strain rate (long-time) response and diffusivity initially show reduced elastic-like contributions and constraints to motion, followed by a surprising increase to values that are significantly higher (i.e., more elastic-like) than their pre-digestion counterparts. Our results can shed important light on the interplay between topology and rheology at different scales, and highlight the unique heterogeneities and hierarchies that arise in topologically-active solutions due to time-varying contributions of topology, crowding, entanglements, and polymer overlap. These out-of-equilibrium fluids may have applications in timed self-assembly, selective filtration and sequestration, and wound-healing.

## Results

### Enzymatic digestion of circular DNA initially increases elastic contributions to the nonlinear force response followed by a decrease to a more viscous-dominated regime

We perform nonlinear optical tweezers microrheology (OTM) on enzymatically active solutions by pulling a bead a distance *s* = 30 μm through the solution at a constant speed of *v* = 60 μm/s. Following this strain, we hold the bead fixed and measure the force decay as the system relaxes into a new mechanically-steady state. The distance and speed were chosen to both be well above the intrinsic lengthscales and relaxation rates of the system to probe the nonlinear response. Namely, the fully extended length of PYES is *L* ≃ 1.97 μm and the radius of gyration of the supercoiled, ring and linear topologies are *R*_*G,S*_ ≃ 103 nm, *R*_*G,R*_ ≃ 135 nm, and *R*_*G,L*_ ≃ 213 nm ^17^. Moreover, the strain speed can be approximately converted to a strain rate of 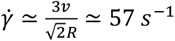 with *R* the bead radius ^38^, with a corresponding timescale 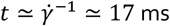, well above the intrinsic Rouse times τ_*R,R*_ ≃ 42 s and τ_*R,L*_ ≃ 136 s for circular and linear PYES chains, respectively ^17,19,39^.

As shown in Figure 2A, the force response to nonlinear straining at all times exhibits an initial increase in force followed by a rolling over to a strain-independent plateau, indicative of a viscous response. The average value of this plateau force *F*_*p*_ increases in the first 10 mins after digestion begins (*t*_*a*_ = 10 mins), followed by large drops from *t*_*a*_ =20 min to *t*_*a*_ = 50 mins, then a more modest decrease before reaching a *t*_*a*_-independent plateau starting at *t*_*a*_ ≈100 mins (Fig 2B). Moreover, the force response for the initial time point (*t*_*a*_ = 10 mins) exhibits a stress overshoot, seen in macrorheology measurements of highly entangled linear polymers ^18,40^, but is typically not seen in microrheology measurements ^41,42^, demonstrating strong entanglements at this point in the digestion process.

**Figure 2:**
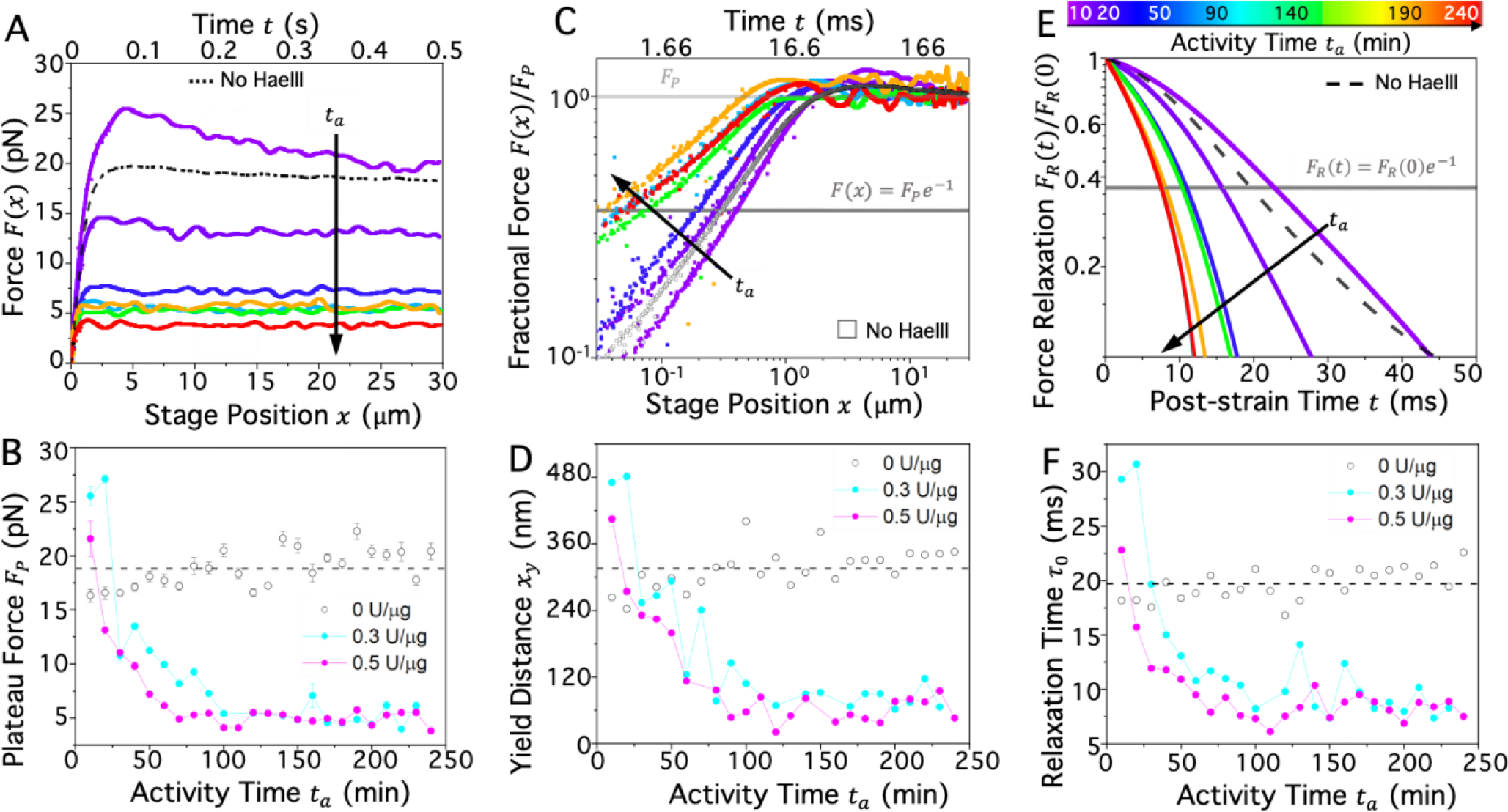
Nonlinear OTM reveals transient increase followed by gradual decrease of resistive force and relaxation timescales of topologically-active circular DNA undergoing linearization and fragmentation. **(A)** Force exerted by the DNA as the trapped probe is moved relative to the sample, by increasing the *x*-position of the stage, for increasing activity times during HaeIII digestion. *t*_*a*_ = 0 denotes the time at which 0.5 U/μg of HaeIII was added to the DNA solution, and colors correspond to different activity times *t*_*a*_according to the color scale in (E). The dashed black curve is the control case in which no enzyme is added. After an initial increase compared to the control, the maximum plateau force *F*_*p*_ monotonically decreases with increasing activity time. **(B)** The time-course of *F*_*p*_ for solutions with 0 (open circles), 0.3 (filled cyan circles) and 0.5 (filled magenta circles) U/μg HaeIIII. Each *F*_*p*_ value plotted is the average force value for *x* = 10 − 30 μ**m** with error bars representing standard error. The time-average of the 0 U/μg data shown is plotted as a dashed line for clarity. The initial increase in *F*_*p*_, as well as the subsequent decrease, is slightly prolonged at lower enzyme concentration (cyan) but the functional form is preserved. **(C)** Force *F* shown in (A), normalized by the corresponding plateau force value *F*_*p*_ (light grey line) and plotted on a log-log scale shows that the distance at which the system yields to the terminal viscous-dominated plateau regime, i.e., the yield distance *x*_*y*_, decreases with increasing time *t*_*a*_. The yield distance is quantified as the displacement at which 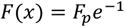 (grey line in plot). **(D)** The dependence of the yield distance on *t*_*a*_ shows similar trends as the plateau force shown in B. **(E)** Force relaxation *F*_*R*_(*t*) measured during the relaxation phase, when the probe is held fixed at *x* = 30 μ**m**, normalized by the value at the beginning of the relaxation phase *F*_*R*_(0). A characteristic relaxation timescale τ_0_ is quantified as the time at which *F*_*R*_(*t*) = *F*_*R*_(0)*e*^−1^ (grey line). **(F)** The trajectories of τ_0_ versus *t*_*a*_ shows similar trends as the plateau force and yield distance shown in B and D.

The general trend of initially increasing in force then decreasing aligns with the results of Ref ^43^ if we attribute the initial rise to the linearization of rings, which increases polymer overlap and entanglements, followed by the drop that was previously seen for viscosity of linear chains undergoing fragmentation. Moreover, the functional form of the decay and the final value of the plateau force is relatively insensitive to enzyme:DNA stoichiometry, with the lower value (0.3 U/μg) reaching a higher initial plateau force *F*_*p*_(*t*_*a*_ = 10 min) and reaching the terminal plateau value *F*_*p*_ slightly later in time (∼100 mins vs ∼75 mins). This dependence aligns with a modestly slower digestion rate, which is predicted to scale with stoichiometry, for 0.3 U/μg compared to 0.5 U/μg, but with the same mechanisms driving the rheological changes.

We also examine the small-strain behavior, during the initial elastic-like increase in force (0 < ***x*** < 1 μm), to determine how the elastic response varies during digestion (Fig 2C). For all activity times, the initial force regime displays approximate powerlaw scaling with strain distance *x, F*(*x*)∼*x*^*b*^ with the scaling exponents *b* generally decreasing with increasing activity time. This effective softening with increasing *t*_*a*_ can be quantified by computing the yield distance *x*_*y*_ at which the force reaches *F*(*x*) = *F*_*p*_*/e*, signifying yielding from elastic-like to viscous-dominated response. Rescaling each force trace by its corresponding plateau value *F*_*p*_, clearly shows the decrease in yield distance, denoted by the *x* value at which each *F*(*x*)*/F*_*p*_ trace crosses the grey horizontal line in Fig 2C, over the course of digestion. Moreover, the trajectory of *x*_*y*_ as a function of activity time (Fig 2D) is highly similar to that of the plateau force (Fig 2B), with an initial rise in *x*_*y*_ followed by a sharp drop before reaching a terminal minimum value at ∼100 min.

We expect this yield distance to be related to a characteristic confinement lengthscale, e.g., the tube diameter *d*_*T*_ for entangled polymers ^19^, that sets the elastic contribution to the response. Namely, the motion of each polymer is constrained by steric interactions with the surrounding polymers, forcing it to undergo elastic-like entropic stretching, rather than dissipative flow, in response to strain. Once the polymer has been strained a distance several times beyond this confinement lengthscale it can no longer maintain its entanglements with the surrounding polymers and instead the strain induces dissipative polymer disentanglement and flow ^18^. Theory describing the dynamics of entangled rings have predicted that the tube diameter for entangled ring polymers is smaller than its linear counterpart via the relation *d*_*T,R*_ ≈0.7*d*_*T,L*_ ^44^, in line with previously reported relation for 115 kbp DNA of *d*_*T,R*_ ≈0.75*d*_*T,L*_ ^39^, suggesting that *x*_*y*_ should be proportionally larger when the solutions are primarily linear versus circular. Importantly, this description is only valid for systems with ample entanglements or other steric interactions, whereas unentangled polymers exhibit a very limited elastic regime (small *x*_*y*_) as the chains can more readily flow in response to strain.

The trajectory of *x*_*y*_ is qualitatively in line with this physical picture, as well as our interpretation of the plateau force data, as we expect initial linearization of the circular DNA to increase *x*_*y*_, while fragmentation, which breaks and removes entanglements, to decrease *x*_*y*_ (Fig 2D). Moreover, the maximum yield distance we measure, which we interpret to be when the polymers are mostly linear, is *x*_*y,L*_ ≃ 480 nm, compared to *x*_*y*,0_ ≃ 315 nm for the initial solution of circular polymers. The ratio *x*_*y*,0_*/x*_*y,L*_ ≃ 0.66 is quite close to the theoretical and experimental ratios *d*_*T,R*_*/d*_*T,L*_ discussed above, corroborating the concept that the yield distance is set by the tube diameter. Moreover, we can compute a tube diameter for linear DNA via *d*_*T,L*_ ≈(*l*_*k*_*l*_*e*_)^1*/*2^ where *l*_*k*_ ≃ 100 nm is the Kuhn length, *l*_*e*_ ≈*l*_*k*_(*c/c*^*^)^1*/*2^ is the entanglement length, and *c*^*^ is the value for purely linear DNA ^18,45^, providing *d*_*T,L*_ ≈160 nm and *x*_*y,L*_ ≈3*d*_*T,L*_. This relation quantitatively shows that yielding occurs once the polymers are strained beyond *x* ≈3*d*_*T*_, a reasonable assumption based on the physical picture described above. It follows that the ring polymers exhibit the same yielding behavior, with *d*_*T*,0_ ≈110 nm and *x* ≈3*d*_*T*_. Finally, the large drop in *x*_*y*_ as digestion proceeds is consistent with fragmentation that reduces and eventually eliminates entanglements.

Next, we turn to the relaxation of the force following straining to determine how the longest relaxation timescales vary during digestion. As shown in Figure 2E, the force relaxation follows similar trends as the response during strain, with the decay in *F*_*R*_ first becoming slower, seen as a shallower slope for *t*_*a*_ = 10 min, followed by rapidly increasing decay rates, seen as steeper slopes that approach a relative constant value for *t*_*a*_ > 50 mins. To approximate a single characteristic relaxation time τ_0_ for the system at each point during digestion, we compute the measurement time at which *F*_*R*_(*t*) = *F*_*R*_(0)*/e* (Fig 2E). As shown in Figure 2F, τ_0_ exhibits the same dependence on activity time as *F*_*p*_ and *x*_*y*_, corroborating the physical picture that the elastic contribution to the nonlinear stress response initially increases, due to linearization of the DNA circles, then yields to a viscous-dominated regime dictated by the fragmentation of the linear chains.

To shed light on the principal mechanism underlying the force relaxation, we note that previous studies examining the nonlinear response of entangled DNA solutions have shown that the relaxation modes that generally contribute to the force relaxation are elastic Rouse relaxation, which occurs over the Rouse time 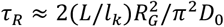 where *D*_0_ is the dilute limit diffusion coefficient; reptation, which occurs over the disengagement time τ_*D*_ ≈3(*L/l*_*e*_)τ_*R*_; and entanglement, which occurs over the entanglement time τ_*e*_ ≈(*l*_*e*_*/L*)^2^τ_*R*_ ^18,30,39,46–48^. For linear DNA solutions, these predicted expressions yield τ_*R*_ ≈136 ms, τ_*D*_ ≈3245 ms, and τ_*e*_ ≈2 ms. For rings we compute, τ_*R*_ ≈42 ms and τ_*e*_ ≈20 ms using the expressions above, and for τ_*D*_, we use the predicted and experimentally validated relation τ_*D,R*_ ≈0.3τ_*D,L*_ to compute τ_*D*_ ≈973 ms ^39,44^. Finally, for supercoiled DNA, τ_*R*_ ≈19 ms, but there are no equivalent τ_*e*_ or τ_*D*_ values, as the concentration for purely supercoiled molecules is 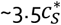 (versus 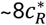 and 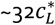 for rings and linear DNA) which is below the nominal entanglement concentration of ∼6*c*^*^. Considering these values, it is clear that reptation is not a primary mode of nonlinear stress relaxation, as τ_*D,R*_ and τ_*D,L*_ are ∼1-2 orders of magnitude slower than τ_0_ ≈6 − 30 ms (Fig 2F). This assumption is also consistent with our finding that all solutions reach a terminal viscous plateau, and with previous studies that have shown that in certain cases even well-entangled linear DNA subject to nonlinear strains can relax their stress without reptation ^42^. Our measured τ_0_ values are, however, comparable to both τ_*R*_(≃ 19, 42, 136 ms) and τ_*e*_(≃ 2, 20 ms) for the different topologies, so we suggest that relaxation proceeds via a combination of both relaxation modes, aligning with previous studies that have shown that nonlinear force relaxation for entangled DNA is typically described by 2-3 modes ^30,39^. Given the topological complexity and polydispersity of our solutions during digestion, we do not expect exact agreement with predictions for monodisperse systems, but order of magnitude agreement with predictions serve as a guide towards underlying mechanisms.

### Linear microrheological response shows opposite dependence on activity time at low frequencies versus high frequencies

To determine how topological digestion of the DNA solutions alters the linear viscoelastic response, we measure the thermal fluctuations of a trapped probe to extract the frequency-dependent linear elastic and viscous moduli ***G***′(***ω***) and ***G***′′(***ω***) (Fig S3). These measurements probe smaller lengthscales and slower strain rates than nonlinear measurements, with a range of accessible frequencies of ***ω*** = **1** − **300** rad s^-1^, corresponding to timescales of ***t*** = **2*π ω***^−**1**^ = 21 ms – 6.25 s, which are all longer than the nominal nonlinear strain timescale 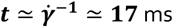. To examine the elastic contribution to the linear stress response we evaluate ***g***^′^(***ω***) = ***G***′(***ω***)*/*|***G***^*^(***ω***)| where |***G***^*^(***ω***)| = [(***G***′(***ω***))^**2**^ + (***G***′′(***ω***))^**2**^]^**1***/***2**^ is the magnitude of the complex modulus, and ***g***^′^(***ω***) ranges between 1 for a purely elastic response to 0 for purely viscous response. Figure 3A reveals distinct dependence on the activity time at high frequencies versus low frequencies. Specifically, for ***ω*** = **30** − **300** rad s^-1^, ***g***^′^(***ω***) exhibits a similar dependence on ***t***_***a***_ as the nonlinear response metrics, with ***g***^′^ generally decreasing with increasing ***t***_***a***_. However, for much lower frequencies of ***ω*** = **1** − **5** rad s^-1^, we observe a counterintuitive nearly opposite trend, with ***g***^′^(***ω***) first decreasing below that of the control, followed by an increase, suggestive of increased elasticity upon fragmentation.

**Figure 3:**
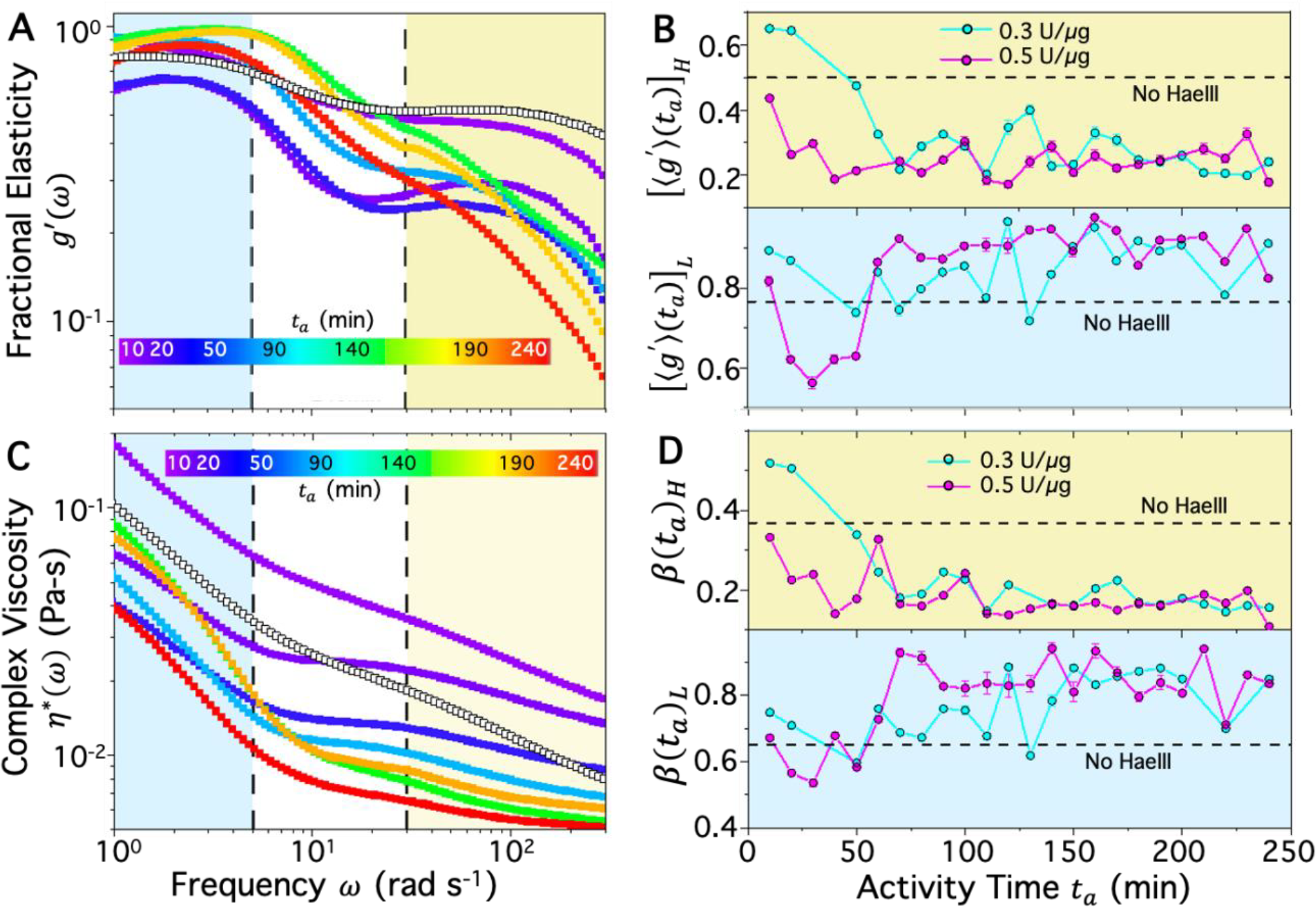
Linear OTM reveals distinct trajectories of viscoelastic moduli at high and low frequencies during topological digestion. **(A)** Fractional elasticity 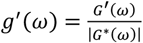versus frequency *ω* for increasing activity times *t*_*a*_ after adding 0.5 U/μg of HaeIII, denoted by the color scale. Control data (0 U/μg) is shown as open symbols. The dependence of *g*^′^(*ω*) on activity time *t*_*a*_ differs at low frequencies (blue, *ω* = 1-5 rad/s) compared to the high frequency regime (yellow, *ω* = 30-300 rad/s). (B) Trajectories of *g*^′^, averaged over all frequencies in the high frequency regime [**⟨***g*^′^**⟩**(*t*_*a*_)]_*H*_ (top, yellow) and low frequency regime [**⟨***g*^′^**⟩**(*t*_*a*_)]_*L*_ (bottom, blue) for HaeIIII:DNA stoichiometries of 0.3 U/μg (filled cyan circles) and 0.5 (filled magenta circles) U/μg, compared to the average value for the undigested control case (dashed line). (C) Complex viscosity *η*^*^(*ω*) versus frequency *ω* for increasing activity times *t*_*a*_ after adding 0.5 U/μg of HaeIII, denoted by the color scale. Control data (0 U/μg) is shown as open symbols. The power-law scaling *η*^*^(*ω*)∼*ω*^−*β*^ is generally steeper (larger *β*) at low frequencies (blue region) compared to high frequencies (yellow region), and depends on *t*_*a*_. (D) Trajectories of *β*(*t*_*a*_)_*H*_ (top, yellow) and *β*(*t*_*a*_)_*H*_ (bottom, blue) determined via fits of *η*^*^(*ω*) to the high and low frequency ranges, respectively, at each activity time *t*_*a*_ for HaeIIII:DNA stoichiometries of 0.3 U/μg (filled cyan circles) and 0.5 (filled magenta circles) U/μg, compared to the average value for the undigested control case (dashed line). At high frequencies the time dependence matches that observed in the nonlinear regime, while a nearly opposite trend is seen at low frequencies.

By averaging ***g***^′^(***ω***) over all frequencies in each regime, we can more clearly compare trajectories of ***g***^′^ in the high and low frequency ranges, [**⟨*g***^′^**⟩**(***t***_***a***_)]_***H***_ and [**⟨*g***^′^**⟩**(***t***_***a***_)]_***L***_ (Fig 3B). The difference from the control data is less extreme compared to the nonlinear regime metrics (Fig 2), but the different trends are obvious. [**⟨*g***^′^**⟩**(***t***_***a***_)]_***H***_ follows a similar trajectory as the nonlinear regime data, reaching a ***t***_***a***_-independent plateau that is lower than the control, whereas the terminal ***t***_***a***_-independent plateau in [**⟨*g***^′^**⟩**(***t***_***a***_)]_***L***_ is higher than the control. The takeaway is that, at long times, the solutions appear to be locally more entangled and elastic-like following digestion, whereas, at short times, the local response is suggestive of fewer entanglements and elastic-like contributions.

We also evaluate the complex viscosity ***η***^*^(***ω***) = |***G***^*^(***ω***)| ***ω***^−**1**^ which shows shear-thinning ***η***^*^(***ω***)∼***ω***^−***β***^ for all frequencies and activity times. For highly entangled linear polymers, ***β*** ≈**1** ^**19**^, whereas rings typically display lower exponents of ***β*** ≈**0**. 6 ^**30**^ and unentangled semidilute polymers display ***β*** ≈**0. 5** ^**49**^. Figure 3c reveals that the thinning exponents are generally higher in the low frequency range (***ω*** = **1** − **5** rad s^-1^) versus the high frequency regime (***ω*** = **30** − **300** rad s^-1^), indicative of stronger constraints. More importantly, the trajectory of ***β*** with activity time displays the same switch between high and low frequency regimes. Namely, at low frequencies, ***β***(***t***_***a***_)_***L***_ generally increases with activity time, similar to [**⟨*g***^′^**⟩**(***t***_***a***_)]_***L***_, reaching a plateau value that is higher than that of the control. Conversely, ***β***(***t***_***a***_)_***H***_ generally decreases with increasing ***t***_***a***_, with the trajectory mirroring [**⟨*g***^′^**⟩**(***t***_***a***_)]_***H***_, reaching a plateau that has a lower value than the control. These results indicate that the local dynamics are dominated by different mechanisms at short times ***t*** = **2*π ω***^−**1**^ of 21 ms ≲ ***t***_***s***_ ≲ 210 ms versus long times 1.25 s ≲ ***t***_***l***_ ≲ 6.3 s.

To consider the differences between these timescales relative to the intrinsic relaxation timescales of the system, we recall the predicted topology-dependent values for ***τ***_***D***_, ***τ***_***R***_ and ***τ***_***e***_ discussed in the previous section. The long-time regime extends to times longer than all of the predicted relaxation timescales, with ***τ***_***D***,***L***_ ≃ **3. 2 *s*** falling within the long-time range and ***τ***_***D***,***R***_ ≃ **0**. 9**7 *s*** lying just short of ***t***_***s***_, so we expect reptation of the long chains to dominate the response. Conversely, the short-time regime is below ***τ***_***D***,***L***_ and ***τ***_***D***,***R***_, as well as above ***τ***_***e***,***L***_ and ***τ***_***e***,***R***_, suggesting that unentangled Rouse dynamics likely dictate the stress response and that the primary contributors to this short-time stress response are ostensibly unentangled circular DNA and shorter linear fragments that have terminal relaxation times much faster than ***τ***_***D***,***L***_ and ***τ***_***D***,***R***_. We will consider this argument further in the Discussion.

### DNA transport exhibits counterintuitive slowing during enzymatic fragmentation

To shed light on the timescale-dependent impact of topological alteration on rheology, we turn to measuring how digestion alters the apparent diffusion of DNA molecules in the topologically-active solutions at long timescales. We use DDM for this purpose, rather than single-molecule tracking, to allow for more statistics in the long-time regime (Figs 1G-J, 4). As shown in Figure 4A, DDM is able to extract statistically meaningful intermediate scattering functions (ISFs) for lag times up to ***t*** = 10 s. Visual inspection of the ISFs for different activity times ***t***_***a***_ (Fig 4A) reveals that ISFs decay more slowly as ***t***_***a***_ increases, indicative of slowing down of dynamics during digestion, in qualitative alignment with the low-frequency microrheology data (Fig 3). By fitting the ISFs for each wave vector ***q*** = **2*π λ***^−**1**^ to an exponential (see Methods) we extract the characteristic decay time ***τ***(***q***) for each time ***t***_***a***_, which is a measure of how quickly density fluctuations at a lengthscale ***λ*** become decorrelated ^**50–52**^. Increasing ***τ***(***q***) values at a given ***q*** indicate slowing of dynamics, which Fig 4B shows occurs as ***t***_***a***_ increases. Moreover all ***τ***(***q***) data roughly follow power-law scaling ***τ***(***q***)∼***q***^−**2**^, expected for diffusive dynamics (rather than subdiffusive or superdiffusive) where the diffusion coefficient is computed via ***D*** = ***τ***^−**1**^***q***^−**2 33**,**51**,**53**,**54**^

**Figure 4:**
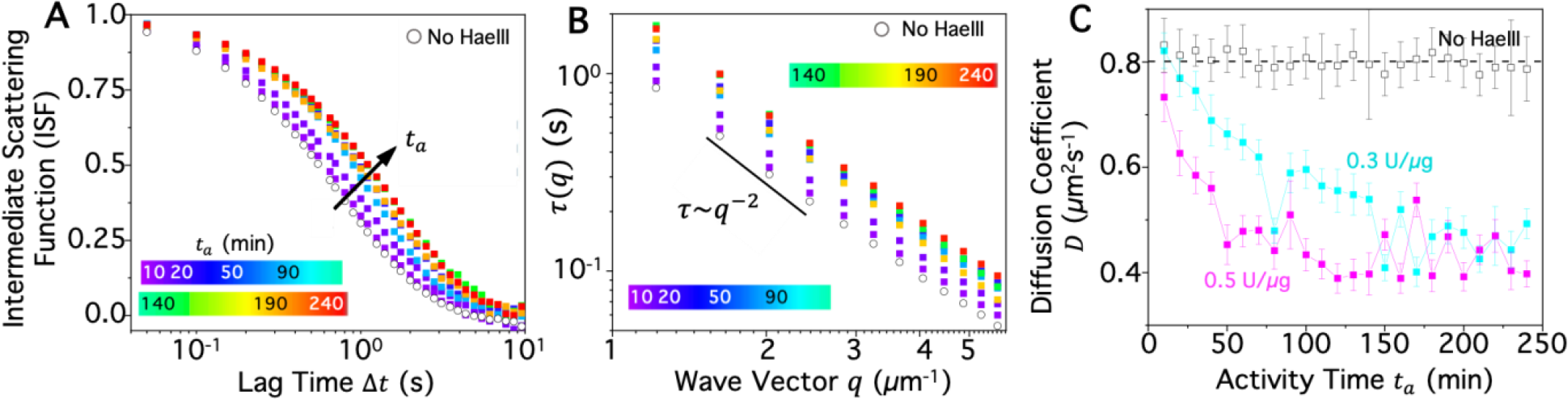
DDM of fluorescent-labeled DNA shows that HaeIII-mediated fragmentation of DNA surprisingly slows the diffusion of DNA. **(A)** Intermediate scattering function (ISF) of topologically-active DNA solution as a function of lag time Δ*t* at different activity times *t*_*a*_ during digestion by 0.5U /μg HaeIII, color coded according to the legend. As digestion proceeds (*t*_*a*_ increases) the time at which ISFs decay is extended, indicating a slowing of dynamics. **(B)** Decay time τ(*q*) vs wave vector *q*, computed via fitting the ISF, shows diffusive dynamics for all times *t*_*a*_, i.e., τ(*q*)∼*q*^−2^ but the value of τ(*q*) at each *q* increases with increasing *t*_*a*_, indicating lower diffusion coefficients *D* (slower dynamics) which can be computed via the relation τ(*q*) = 1*/Dq*^2^. **(C)** Diffusion coefficients *D*, determined from τ(*q*) versus activity time for HaeIII:DNA stoichiometries of 0.3 U/μg (filled cyan circles), 0.5 U/μg (filled magenta circles) and 0 (open circles, dashed line denoted average).

Figure 4C plots ***D*** versus ***t***_***a***_, as determined from fits of ***τ***(***q***), revealing a strikingly similar trajectory as [**⟨*g***^′^**⟩**(***t***_***a***_)]_***L***_ and ***β***(***t***_***a***_)_***L***_, with an initial slowing of dynamics (expected for increased elasticity and entanglements) until reaching a plateau at ***t***_***a***_ ≈**100** s for 0.5 U/μg. The lower stoichiometry (0.3 U/μg) decays more slowly due to slower digestion, as seen in the OTM data, albeit to a lesser extent.

To determine how general this result is, we perform DDM on solutions undergoing digestion by another enzyme, ApoI, which cuts the DNA at a different number of recognition sites compared to HaeIII (8 vs 17) and the resulting fragments have sticky overhangs rather than the blunt ends that HaeIII cleaving provides (Fig S2). As shown in Figure 5A, ***τ***(***q***) increases with increasing activity time, just as we see for HaeIII (Fig 4B), and in fact follows a trajectory closer the that for 0.5 U/μg HaeIII. To ensure that this slowing is not an artefact of our DDM analysis, we also track single DNA molecules in the topologically-active solutions and evaluate their mean-squared displacements (MSD) versus lag time. Figure 5B shows that indeed the diffusive slowing is preserved in the tracking data, with the MSDs decreasing with increasing activity time, indicative of slower dynamics, and the motion is primarily diffusive, as indicated by the linear scaling of the MSDs with lag time (i.e., ***MSD***∼***t***^***α***^ with ***α*** ≃ **1**). From linear fits to the MSDs, we compute the diffusion coefficients that correlate remarkably well with those measured via DDM over the full time-course of activity (Fig 5C). For different measurement techniques, enzymes and stoichiometries, the slowing of DNA dynamics is universal and trajectories are largely insensitive to the details.

**Figure 5:**
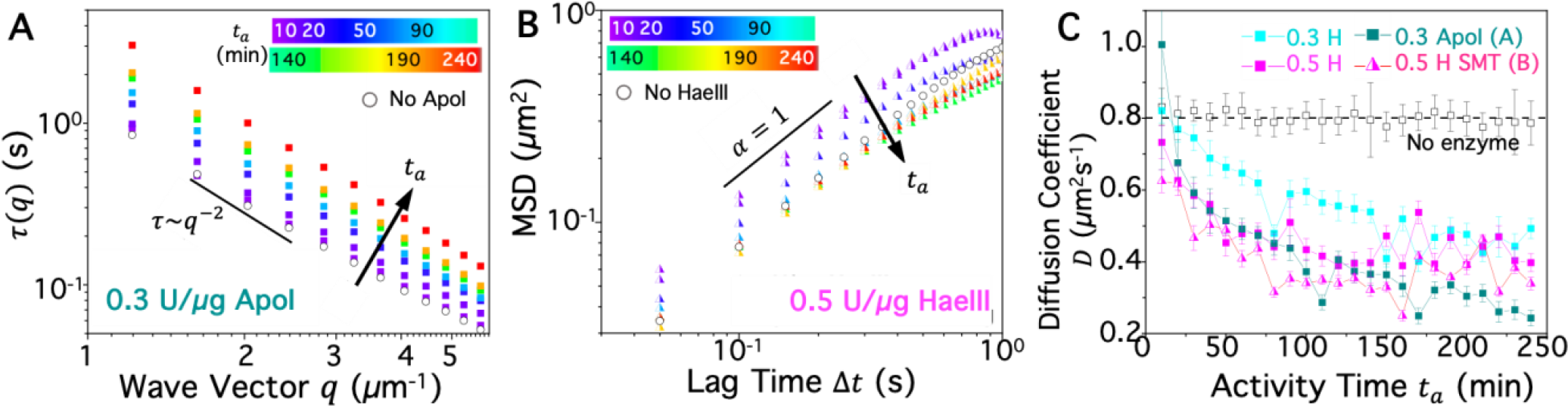
The apparent slowing of DNA transport observed at long times is universal across analysis methods as well as enzyme type and concentration. **(A)** Decay time τ(*q*) vs wave vector, *q* computed identically as in Fig 4B, for DNA solutions undergoing digestion by the multi-site restriction endonuclease, ApoI, at a ApoI:DNA stoichiometry of 0.3U/μg. ApoI digestion produces 8 fragments with sticky overhangs, compared to 17 blunt-ended fragments produced by HaeIII. The observed *q*-scaling of τ(*q*) and its increase with increasing *t*_*a*_ are very similar to those seen in Fig 4B for 0.5 U/μg HaeIII. **(B)** Tracking of single fluorescent-labeled DNA molecules in the same conditions shown in Fig 4A,B (0.5 U/μg HaeIII) shows that the mean-squared displacement (*MSD*) of the molecules are largely diffusive, i.e., *MSD*∼Δ*t*^α^ with α = 1, similar to results from DDM. Morevoer, the magnitude of the *MSD* at any given lag time Δ*t* decreases with increasing activity time *t*_*a*_, indicating a slowing of molecular diffusion, similar to that seen in A and Fig 4. **(C)** Diffusion coefficients *D* calculated using DDM and single-molecule tracking (SMT) for varying concentrations (0.3 U/μg (cyan, teal), 0.3 U/μg (magenta, pink)) and types (HaeIII (H), ApoI) of enzymes show universal slowing of diffusivity during digestion that is rapid initially and then weaker, appearing to asymptote to a steady-state value when digestion is complete. ApoI and SMT data are determined from A and B while HaeIII DDM data are the same as in Fig 4C.

**Figure 6:**
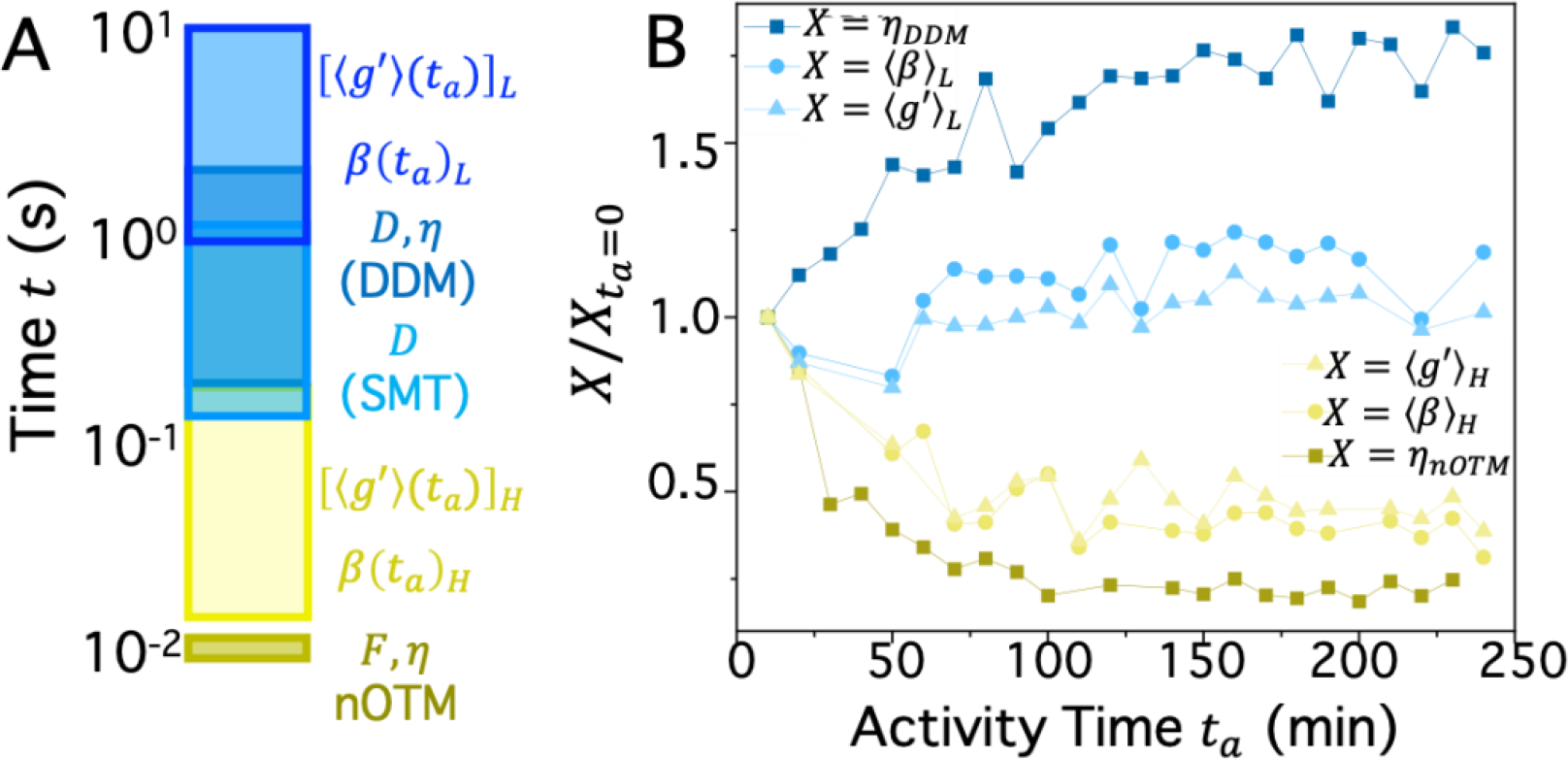
Different relaxation mechanisms manifest at short and long timescales during digestion of circular DNA into polydisperse fragments. (A) Three decades of timescales that different measurement techniques and metrics probe, colorized according to short (yellow shades) and long (blue shades) timescales. Low-frequency linear OTM, DDM, and SMT probe long timescales (∼0.2 – 6 s) while high-frequency linear OTM and nonlinear OTM probe fast timescales (∼17-200 ms).Various metrics of confinement *X*, shown in previous figures, normalized by their *t*_*a*_ = 0 values 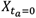, and averaged over the 0.3 and 0.5 U/μg HaeIII conditions, are color coded according to the timescale that the measurement probes (short/yellow or long/blue). The different symbols and shades of each color indicate the metric or measurement technique according to the legend. For DDM and nonlinear OTM, the normalized effective viscosities 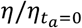 are plotted, computed as 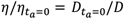 and 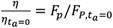, respectively. Short timescale metrics universally indicate that topological activity increases dissipation and mobility, while the opposite is observed at long timescales.

## Discussion

The data discussed above shows stark differences between how linearization and fragmentation alter the rheology and dynamics at different timescales. The short-time dynamics make intuitive sense. Namely, as the DNA gets chopped into smaller and smaller pieces the elastic contribution to the stress response goes down. However, upon initial linearization of the rings this elastic contribution increases as linear chains form stronger entanglements than rings and the degree of polymer overlap at a given concentration is likewise higher. These general trends were also observed in recent particle-tracking microrheology experiments that showed that linearization of circular 5.9-kbp DNA lead to a steady increase in viscosity, while fragmentation of linear 48.5-kpb DNA led to a steady decrease ^**13**^.

To understand the long-time behavior, we turn to recent work, described in the Introduction, that examined the bulk rheology of topologically-active solutions of 11-kbp circular DNA and their composites with dextran ^**14**^. In these studies, measurements were performed at ***ω*** = **1** rad s^-1^, which is the lowest frequency (longest time) that we evaluate here, over the course of digestion by a multi-site cutting enzyme that cleaved the rings into 13 fragments of different lengths. These experiments showed that fragmentation actually led to an abrupt increase in ***G***′ which only occurred following complete digestion (rather than initial linearization). Similar abrupt jumps in ***G***′ were observed in composites with dextran, but the onset was much earlier in the digestion process. This early onset was shown to be due to crowding of the DNA by dextran that facilitated clustering of the DNA, via depletion interactions, that increased the local polymer overlap and entanglements within the clusters. These ‘slow’ clusters, where the DNA is more constrained and diffuses more slowly, dominate the system dynamics at long times, but are frozen out at short times, below the relaxation timescales of the clustered DNA.

Similar effects of DNA crowding by dextran have been seen in bulk rheology measurements of steady-state DNA-dextran composites (no digestion) ^**55**^. In the absence of dextran, clustering of longer DNA chains is still expected to occur via crowding by the smaller fragments that have similar radius of gyration as dextran ^**14**^. Specifically, enhanced self-association and clustering of molecules in the presence of crowders occurs as the crowding population seeks to maximize its entropy by maximizing its available volume, driving larger molecules together. This depletion effect only occurs, i.e., is only entropically favorable, when the crowders are sufficiently small and at high enough concentration, as compared to molecules being crowded. For reference, the radius of gyration of the dextran used in the previous studies was ***R***_***G***_ ≃ 19 nm while for DNA it was ***R***_***G***_ ≃ 280 nm and ***R***_***G***_ ≃ 196 nm. Here, ***R***_***G***_ ≃103-213 nm for DNA (depending on the topology), while the radius of gyration for the average fragment length at complete digestion, **⟨*l***_***f***_**⟩** ≃ **115** nm, is ***R***_***G***_ ≃ 20 nm and for the smallest length fragment ***l***_***f***,***min***_ ≃ 6.3 nm (19 basepairs) it is ***R***_***G***_ ≃ 5 nm. Given the similar sizes of dextran and the DNA fragments in our current work and previous studies, it is reasonable to assume that similar depletion interactions may be at play. Moreover, the roughly order of magnitude difference between the size of DNA and its crowders (fragments or dextran) in all cases, suggests that we may expect similar mechanisms to drive the non-equilibrium rheological behavior.

Based on this knowledge, we argue that the increased elasticity and slowed diffusion that fragmentation elicits at long timescales is a result of the depletion-driven formation of local clusters of long DNA via crowding by shorter fragments. The DNA molecules within these clusters are more entangled and slower than those in the surrounding solution, and it is the dynamics of these slow clusters that dominate the rheology and diffusion of the system at long times, i.e., those comparable to the relaxation of the clustered DNA. This slowing is not seen at short times as the relaxation of these slow molecules is frozen out on this fast measurement timescale. To corroborate this physical picture, we point to the larger relative drop in DNA diffusion coefficients, measured via DDM and single-molecule tracking, compared to the relative increase in **⟨*g***′**⟩** and **⟨*β*⟩** measured via OTM (Fig 6A). The former measurements rely on fluorescent-labeled DNA to report dynamics while the latter does not. However, DNA below ∼2 kbp cannot be seen in typical fluorescence microscopes due to resolution limits and the inability to add enough fluorophores to smaller chains. Nevertheless, we were still able to visualize labeled DNA throughout the duration of digestion (Fig 1G), suggesting that the DNA that we were tracking were the longest fragments, while the shorter ones were unable to contribute to the signal. Thus, we expect our DDM and tracking data to be representative of only the slow-moving clusters of longer fragments. OTM measurements on the other hand measure the local rheology in the vicinity of the trapped bead, which could be in a cluster or in the faster surroundings, so the apparent increase in elasticity is more subdued.

To more directly compare the changes in rheological properties during digestion, measured over different timescales and with different techniques (Fig 6A), we plot together the trajectories of all metrics shown in Figs 2-4, normalized by their ***t***_***a***_ = **0** value, and averaged over the two stoichiometries (Fig 6B). For more straightforward comparison of metrics that increase with increased elasticity and slowing of dynamics, we convert our DDM-measured normalized diffusion coefficients 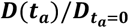 to viscosity via the relation ***D***∼***η***^−**1**^, and our maximum plateau force 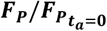 to viscosity via the relation ***F***_***P***_ = ***ηv***. Figure 6B shows the trajectories of these long-time (DDM) and short-time (nonlinear OTM) viscosities, along with **⟨*g***′**⟩** and **⟨*β*⟩** for both short and long measurement timescales. The short-time trajectories of 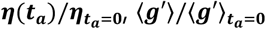, and 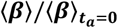, are similar for all three metrics, with the nonlinear OTM viscosity decaying to a slightly lower value, presumably because of the faster timescales probed (∼18 ms vs 21-210 ms). The long-time trajectories show a more pronounced difference, with 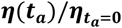 increasing by ∼80% over the course of digestion, while 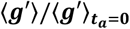, and **⟨**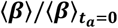 increase by ∼22%. This significant difference corroborates the effect described above of only being able to visualize the larger fragments in DDM (as well as single-molecule tracking, Fig 5B).

## Conclusion

In conclusion, we have presented the results of comprehensive measurements of concentrated circular DNA pushed out-of-equilibrium via the action of multi-site cleaving enzymes that linearize and chop up DNA circles. Using linear and nonlinear optical tweezers microrheology, as well as DDM and SMT, we span over 3 orders of magnitude in measurement timescales to reveal a surprising timescale-dependent variation of viscoelastic moduli and DNA diffusivity during enzymatic activity. Namely, at short timescales, well below the nominal disengagement times for full-length circular and linear DNA, probed via nonlinear OTM (Fig 2) and high-frequency linear OTM (Fig 3), digestion leads to an initial increase in metrics of elasticity and confinement, i.e., *g*′, *β, η, x*_*y*_, τ_0_, due to linearization of the circular DNA, followed by a rapid decrease as fragmentation continues, due to disruption of entanglements and decreased polymer overlap, after which a plateau is reached once complete digestion is achieved. At long timescales, comparable to or longer than the disengagement times for the full-length DNA, probed via low-frequency OTM (Fig 3), DDM (Fig 4) and SMT (Fig 5), we observe remarkably opposite trajectories of metrics, whereby fragmentation causes a rapid increase in elasticity metrics, i.e., *g*′, *η, β*, after an initial transient drop, that stabilizes to a plateau value at nearly the same activity time as the short-time metrics.

We rationalize these results as arising from the different relaxation mechanisms and molecular lengthscales that the two regimes can detect, along with crowding induced self-association of longer DNA chains by short fragments. At short timescales, the relaxation modes of the long DNA strands are prohibitively slow to be detected and are thus frozen out, leaving the shorter fragments and more weakly entangled molecules to dominate the signal. As fragmentation proceeds, these shorter chains become shorter, which decreases overlap and entanglements that provide elasticity. Moreover, the depletion of the longer chains by the shorter chains increases the volume available to the shorter chains, making them effectively more dilute and mobile, further decreasing any elastic contributions to the stress response. Conversely, at long lengthscales, the motion of the slow-moving clusters of long DNA dominates the response. As fragmentation proceeds, the depletion interaction strength increases as more and more fragments are created, increases the self-association of longer chains, which in turn increases their entanglements and constraints and slows their dynamics, leading to an increase in elasticity metrics and slowing of diffusion.

Our results highlight the ability to use topology to drive non-equilibrium restructuring and viscoelastic alterations that can be tuned at the molecular level to elicit starkly different trajectories at different spatiotemporal scales. These hierarchical reconfigurable materials may prove useful in applications in sequestration, self-healing, filtration, and micro-actuation. More generally, the use of enzymes to impart activity into steady-state systems offers a mechanism for self-assembly, reconfigurability, and self-healing with timing and strength that can be precisely programmed by the stoichiometries and types of enzymes and substrate.

## Methods

### DNA

We prepare solutions of double-stranded DNA, of length ***L*** = **5**. 9 kilobasepairs (kbp), via replication of PYES2 plasmid constructs in Escherichia coli followed by extraction, purification and concentrating as described previously ^**56–58**^. Briefly, cultures of E. coli cells containing PYES2 are grown from frozen glycerol stocks, after which the DNA is extracted from cells via alkaline cell lysis, renatured via treatment with an acidic detergent, precipitated via isopropanol, and resuspended in nanopure deionized water (DI). To remove RNA and proteins the solution is treated with Rnase A followed by phenolchloroform extraction and dialysis. The purified DNA solution is further concentrated via rotary vacuum concentration and stored at 4°**C**.

We use gel electrophoresis and band intensity analysis, employing Life Technologies E-Gel Imager and Gel Quant Express software, to determine a stock DNA concentration of ***c*** =8 mg/mL and a topological distribution of ∼55% relaxed circular DNA and ∼45% supercoiled circular DNA. The radius of gyration for each topology is ***R***_***G***,***R***_ ≃ **135** nm and ***R***_***G***,***R***_ ≃ **103** nm, from which we compute an overlap concentration of 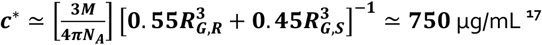

### Restriction Endonucleases

We use the high-fidelity restriction endonuclease HaeIII (New England BioLabs) to introduce topological activity into the DNA solution. HaeIII cleaves pYES2 at 17 different 5’-GG/CC-3’ recognition sites (Figure 1a), converting each circular DNA to linear topology (the first cut) and then cutting the resulting linear strand into 17 blunt-ended linear fragments of different lengths ranging from ***l***_***f***,***min***_ = 19 bp to ***l***_***f***,***max***_ = 1642 bp, with an average fragment length of **⟨*l***_***f***_**⟩** ≃ 345 bp (Fig S1). For all data shown we use a HaeIII:DNA stoichiometry of 0.3 U/μg or 0.5 U/μg, chosen to be low enough to ensure that any topological activity can be considered frozen out on the timescale of each measurement (∼10 s) but fast enough that complete digestion is achieved over the course of each 4 hr experiment. We also performed a subset of experiments using the high-fidelity restriction endonuclease ApoI (New England BioLabs) at stoichiometry of 0.3U/μg DNA. ApoI cleaves pYES2 at 8 different 5’-R/AATTY-3’ recognition sites, resulting in 8 fragments with sticky overhangs and lengths from ***l***_***f***,***min***_ = 11 bp to ***l***_***f***,***max***_ = 2585 bp, with **⟨*l***_***f***_**⟩** ≃ 738 bp (Fig S2).

### Sample preparation

To prepare topologically active DNA solutions for all experiments, we dilute the stock DNA to a final concentration of 6 mg/mL with 1/10^th^ volume of 10× CutSmart Buffer (0.5 M Potassium Acetate, 0.2 M Tris-acetate, 0.1 M Magnesium Acetate; New England BioLabs), 0.1% Tween (to prevent non-specific interactions), and the appropriate volume of HaeIII. We mix the solution by pipetting up and down with a wide-bore pipet tip, and add the enzyme last, noting the time at which it is added, which we define as ***t***_***a***_ = **0**. The buffer conditions and temperature (20°C) provide good solvent conditions for the DNA ^**43**,**45**,**57**,**59**,**60**^.

For DDM and single-molecule tracking, we include 2-20 μg/mL of fluorescent-labelled DNA and an oxygen scavenging system (45 μg/mL glucose, 43 μg/mL glucose oxidase, 7 μg/mL catalase, 5 μg/mL beta-mercaptoethanol) to inhibit photobleaching. The DNA is labelled with covalent dye Mirus-488 (Mirus Bio) at a dye:basepair ratio 1:5 using the Mirus Label IT Nucleic Acid Labeling Kit (Mirus Bio).

For optical tweezers microrheology measurements we add a trace amount of 4.5-μm diameter polystyrene microspheres (Polysciences) that are coated with BSA to inhibit non-specific interactions with the DNA and enzymes.

We load the final 10 μL sample into a microscope sample chamber created by fusing together a glass coverslip and microscope slide with a heated parafilm spacer, and seal the chamber with UV glue. The chamber surfaces are passivated with 10 mg/mL BSA to prevent non-specific adsorption of the DNA or enzyme.

### Optical Tweezers Microrheology

We use optical tweezers microrheology to determine the linear and nonlinear rheological properties over the course of digestion (Fig 1B-F). The optical trap, built around an Olympus IX71 epifluorescence microscope, is formed from a 1064 nm Nd:YAG fiber laser (Manlight) focused with a 60× 1.4 NA objective (Olympus), as described previously ^**49**,**61**^. The force exerted by the DNA on a trapped bead is determined by recording the laser beam deflections via a position sensing detector (Pacific Silicon Sensors) at 20 kHz. The trap is calibrated for force measurement using Stokes drag method ^**41**,**42**,**49**^. All microrheological data is recorded at 20 kHz, and each measurement is taken with a new microsphere in an unperturbed location. We collect data every 10 minutes over the course of 240 mins.

### Linear microrheology

We determine linear viscoelastic properties from the thermal fluctuations of trapped microspheres, measured by recording the associated laser deflections for 100 seconds (Fig 1B,D). We extract the elastic modulus ***G***′(***ω***) and viscous modulus ***G***′′(***ω***), from the thermal fluctuations using the generalized Stokes-Einstein relation (GSER) as described in ref ^**62–64**^. In brief, we compute the normalized mean-squared displacements (***π***(***τ***) = **⟨*r***^**2**^(***τ***)**⟩***/***2⟨*r***^**2**^**⟩**) of the thermal forces, which we convert into the Fourier domain via:

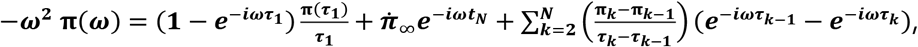

where ***τ***, 1 and ***N*** represent the lag time and the first and last point of the oversampled 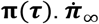 is the extrapolated slope of **π**(***τ***) at infinity. Oversampling is done using the MATLAB function PCHIP. **π**(***ω***) is related to viscoelastic moduli via:

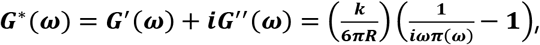

where ***R*** and ***K*** represent the microsphere radius and trap stifness. From ***G***′(***ω***) and ***G***′′(***ω***) (Fig S3), we compute the the magnitude of the complex modulus |***G***^*^(***ω***)| = [(***G***′(***ω***))^**2**^ + (***G***′′(***ω***))^**2 1***/***2**^] and the corresponding fractional elastic modulus ***g***^′^(***ω***) = ***G***′(***ω***)*/*|***G***^*^(***ω***)| (Fig 2A,B) and complex viscosity ***η***^*^(***ω***) = |***G***^*^(***ω***)|*/****ω*** (Fig 2C,D).

#### Nonlinear microrheology

We perform nonlinear microrheology measurements by displacing a trapped microsphere embedded in the sample through a distance ***s*** = **30 *μ*m** at a speed ***v*** = 60 μm/s using a piezoelectric nanopositioning stage (Mad City Laboratories) to move the sample relative to the microsphere (Fig 1B,E), after which we hold the bead fixed for 15 s (Fig 1C,F). We measure the force exerted on the microsphere for the duration of the measurement to determine the force ***F***(***x, t***) the solution exerts to resist the perturbation (Fig 1E), and the relaxation of that force ***F***_***R***_(***t***) as the system relaxes to a new mechanically-steady-state (Fig 1F).

### Differential Dynamic Microscopy (DDM)

We image fluorescent-labeled DNA embedded in each topologically-active DNA solution with an Olympus IX73 epifluorescence microscope with an X-Cite LED light source, 488-nm emission and 535-nm detection filter cube, and a 60x, 1.2NA objective (Olympus). Every 10 minutes after adding the restriction endonuclease ( ***t***_***a***_ = **0**), we capture times-series of images of 1440 x 1920 pixels, providing a field-of-view of 87.3 μm x 116 μm. We use a Hamamatsu Orca-Flash 2.8 to capture images at a frame rate of 20 fps for 80 s. We divide each time-series into 25 separate 256 x 256 pixel regions of interest (ROIs).

We perform DDM on each ROI using the PyDDM package on Github ^**53**^. Following the method of DDM originally described by Cerbino and Trappe ^**51**^, we compute the Fourier transform of the differences between images separated by a given lag time, **Δ*t***, for lag times that range from 50 ms (the time between frames) to 10 s. From this DDM analysis, we obtain the image structure function which we model as ***D***(***q*, Δ*t***) = ***A***(***q***)[**1** − ***f***(***q*, Δ*t***)] + ***B*** where ***A***(***q***) is the amplitude, ***B*** is the background, and ***f***(***q*, Δ*t***) is the intermediate scattering function (ISF). The amplitude and background terms depend on the properties of the imaging system, structure of the sample being imaged, and the camera noise. The ISF contains information about the system dynamics. The ISFs are fit to an exponential function of the form ***f***(***q*, Δ*t***) = **exp**(−(**Δ*t⁄τ***(***q***))^***s***(***q***)^) where ***τ***(***q***) is the characteristic decay time and ***s***(***q***) is a stretching exponent that remained near or just under 1 for our data. For each ROI of the videos recorded every 10 minutes from ***t***_***a***_ = **10** min to ***t***_***a***_ = **2**4**0** min, we determine ***τ***(***q***). For each ***t***_***a***_, we average ***τ***(***q***) across the 25 ROIs to determine the diffusion coefficient according to the equation ***D*** = **1*⁄τq***^**2**^. We compute this diffusion coefficient over the range of ***q*** values from 1 to 2.5 **μm**^−**1**^.

### Single-MoleculeTracking

To track the diffusion of single DNA molecules within topologically-active solutions, we add labeled DNA at a 10x lower concentration compared to samples prepared for DDM analysis. The remaining imaging details and specifications are identical as described above. In every frame ∼200 individual DNA molecules are visible and sufficiently separated and dispersed throughout the field-of-view to allow for accurate single-molecule tracking. We use custom particle tracking scripts (Python) to track the center-of-mass of individual DNA molecules and measure their *x* and *y* displacements (Δ*x*, Δ*y*) for given lagtime Δ*t* ^65,66^. We then compute mean-squared displacements **⟨**(Δ*x*(Δ*t*))^2^**⟩** and **⟨**(Δ*y*(Δ*t*))^2^**⟩** from which we compute an average mean-squared displacement 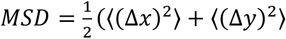as a function of lag time. For MSDs that scale linearly with lag time, i.e., *MSD*∼(Δ*t*)^α^, where α = 1, the diffusion coefficient can be computed via the relation *MSD* = 2*D* Δ*t*. Using this approach, we determine DNA diffusion coefficients by fitting our measured MSDs from Δ*t* = 0.1 s to Δ*t* = 0.7 s, where the data is reliably linear, to a linear function and equating the slope to 2*D*.

## Author Contributions

R.M.R.A. designed the research and supervised experiments. P.N. and R.M.R.A. wrote the paper and interpreted data. P.N. and N.C. conducted experiments and analyzed data. R.M. supervised experiments, and helped with data analysis, interpretation, and writing the paper.

## Acknowledgements

P.N. and R.M. acknowledge the Research Corporation for Scientific Advancement for support. R.M.R.A. acknowledges the US Air Force Office of Scientific Research (Grant no. AFOSR-FA9550-17-1-0249) for support.

