## Supplemental Figures 1-3 for "Enzymatic cleaving of entangled DNA rings drives scale-dependent rheological trajectories"

### Supplemental Information

Figure S1: Digestion map and fragment lengths for HaeIII cleaving of PYES2

Figure S2: Digestion map and fragment lengths for ApoI cleaving of PYES2

Figure S3: Time-resolved linear elastic and viscous moduli, $G'(\omega)$ and $G''(\omega)$, measured during HaeIII activity, from which $g'(\omega)$ and $\eta^*(\omega)$ are computed.

| Fragment lengths (bp) |
| --- |
| 255 |
| 24 |
| 44 |
| 352 |
| 440 |
| 1642 |
| 720 |
| 60 |
| 354 |
| 505 |
| 267 |
| 80 |
| 458 |
| 434 |
| 29 |
| 174 |
| 19 |

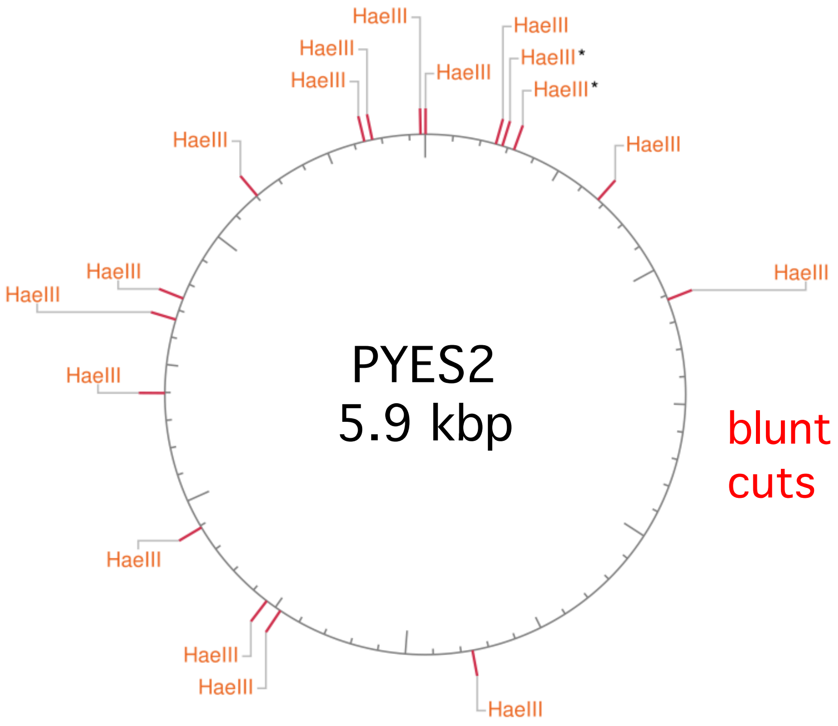


**Figure S1: Digestion map and fragment lengths for HaeIII digestion of PYES2.** HaeIII cleaving sites are indicated by red lines with orange HaeIII labels. The red indicates a blunt cut in which each strand of the double-stranded DNA is cut at exactly the same place. The fragment lengths are listed in order based on the PYES2 sequence which starts at the top of the circular vector map and reads clockwise.

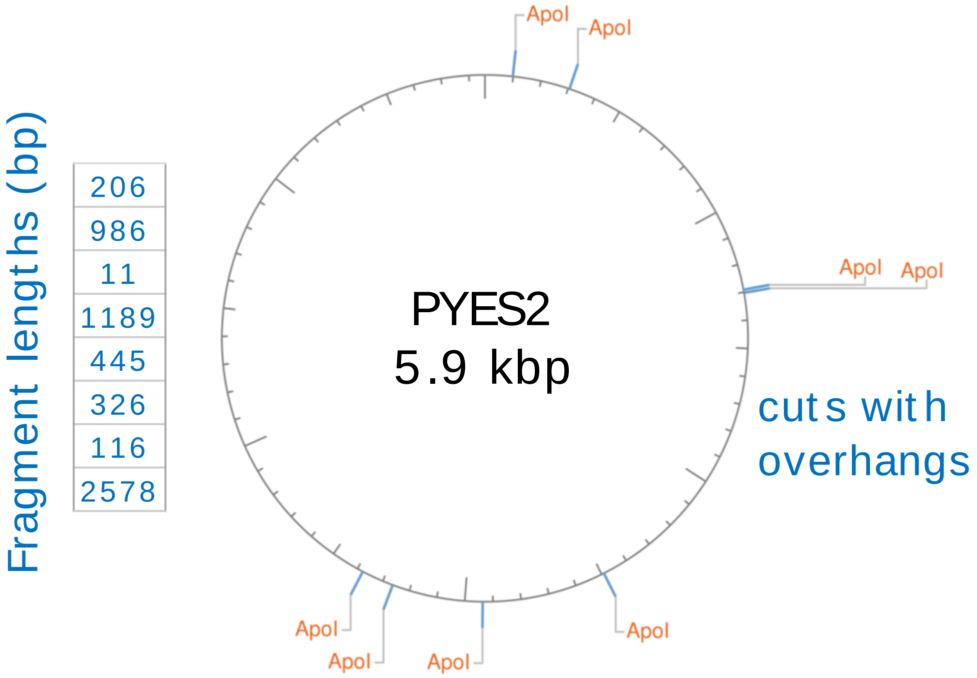


**Figure S2: Digestion map and fragment lengths for ApoI digestion of PYES2.** ApoI cleaving sites are indicated by blue lines with orange ApoI labels. The blue indicates the the enzyme cuts the two strands of the double-stranded DNA at slightly offset locations, resulting in ssDNA overhangs at the ends of the fragment that render them 'sticky'. The fragment lengths are listed in order based on the PYES2 sequence which starts at the top of the circular vector map and reads clockwise.

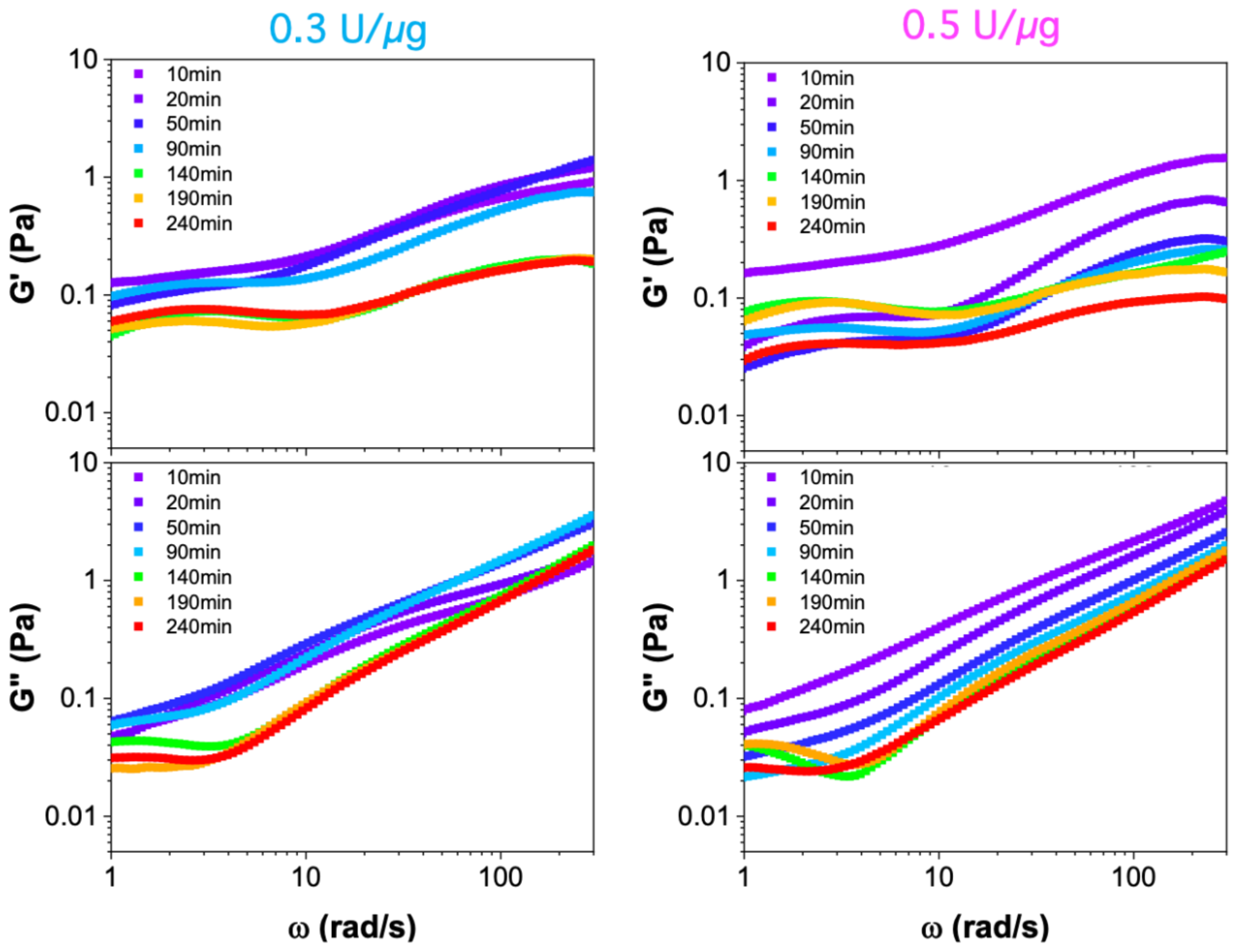


**Figure S3: Time-resolved linear elastic and viscous moduli, $G'(\omega)$ and $G''(\omega)$, measured during HaeIII activity, from which $g'(\omega)$ and $\eta^{*}(\omega)$ are computed.** $G'(\omega)$ (top panels) and $G''(\omega)$ (bottom panels) measured at different activity times ranging from $t_a = 10$ min (purple) to $t_a = 240$ min (red) in solutions with HaeIII:DNA stoichiometries of 0.3 U/μg (left, cyan) and 0.5 U/μg (right, magenta). Data shown is used to compute $g'(\omega)$ and $\eta^{*}(\omega)$ shown in Figure 3 of the main text.
